# Efficient compartmentalization in insect bacteriomes protects symbiotic bacteria from host immune system

**DOI:** 10.1101/2022.03.18.484386

**Authors:** Mariana Galvão Ferrarini, Elisa Dell’Aglio, Agnès Vallier, Séverine Balmand, Carole Vincent-Monégat, Sandrine Hughes, Benjamin Gillet, Nicolas Parisot, Anna Zaidman-Rémy, Cristina Vieira, Abdelaziz Heddi, Rita Rebollo

## Abstract

**Background:** Many insects house symbiotic intracellular bacteria (endosymbionts) that provide them with essential nutrients, thus promoting usage of nutrient-poor habitats. Endosymbiont seclusion within host specialized cells, called bacteriocytes, often organized in a dedicated organ, the bacteriome, is crucial in protecting them from host immune defenses while avoiding chronic host immune activation. Previous evidence obtained in the cereal weevil *Sitophilus oryzae* has shown that bacteriome immunity is activated against invading pathogens, suggesting endosymbionts might be targeted and impacted by immune effectors during an immune challenge. To pinpoint any molecular determinants associated with such challenges, we conducted a dual transcriptomic analysis of *S. oryzae*’s bacteriome subjected to immunogenic peptidoglycan fragments.

**Results:** We show that upon immune challenge the bacteriome actively participates in the innate immune response via an induction of antimicrobial peptides (AMPs). Surprisingly, endosymbionts do not undergo any transcriptomic changes, indicating that this potential threat goes unnoticed. Immunohistochemistry showed that TCT-induced AMPs are located outside the bacteriome, excluding direct contact with the endosymbionts.

**Conclusions:** This work demonstrates that endosymbiont protection during an immune challenge is mainly achieved by efficient confinement within bacteriomes, which provides physical separation between host systemic response and endosymbionts.

## Background

Nutritional symbiosis between animals and microorganisms is a major driver of adaptation [1] as it participates in the colonization of nutrient-poor environments by complementing the metabolic needs of the host [2]. Notably, thanks to intracellular symbiotic bacteria (endosymbionts), insects can thrive on unbalanced carbohydrate-based diets, including blood, plant sap, or cereal grains [1,3–6]. However, the constant presence of microorganisms within an insect’s body represents a permanent challenge for the immune system [7]. The host immune system must conserve its ability to react against pathogens, while keeping beneficial symbionts alive and metabolically active [8]. The establishment of an equilibrium between excessive host colonization by the symbiont and chronic activation of the host immune system is essential in such symbiotic relationships, as the former would be detrimental for host survival, while the latter would result in symbiotic damage and host fitness reduction [9]. To better understand the co-evolution between the host immune system and the intracellular symbiotic bacteria, it is therefore important to pinpoint the molecular determinants of endosymbiont tolerance and pathogen control.

The association between the cereal weevil *Sitophilus oryzae* and its recently-acquired Gram-negative intracellular bacterium, *Sodalis pierantonius* (~28K years, [10,11]), is a remarkable example of homeostasis between insects and endosymbionts. *S. pierantonius* are contained within specialized gigantic cells, the bacteriocytes, which at the larval stages are located in a specialized organ – the bacteriome – at the foregut-midgut junction [3,12]. While wild *S. oryzae* animals are always associated with *S. pierantonius*, comparative studies between symbiotic and artificially-obtained aposymbiotic insects have shown that the presence of the endosymbiont accelerates insect development, allows strengthening of the insect cuticle [13], and enables flying [14].

Contrary to most long-lasting insect endosymbionts, the *S. pierantonius* genome contains genes encoding a functional type III secretion system (T3SS) [15], which was shown to be necessary during insect metamorphosis, where host stem cells are infected by the endosymbiont, followed by bacteriocyte differentiation and adult bacteriome formation [16]. The *S. pierantonius* genome also encodes genes necessary for Microbial-Associated Molecular Patterns (MAMPs) synthesis, including peptidoglycans (PGs), which are able to activate the insect immune responses through their interaction with host pattern recognition receptors [7]. Injection of *S. pierantonius* into the insect hemolymph triggers the production of a plethora of antimicrobial peptides (AMPs) [17], suggesting its presence within the host body is an ongoing immune threat. Nevertheless, chronic immune system activation is avoided by the compartmentalization of the endosymbiont within bacteriocytes and the expression of an adapted local immune system [17–20]. The coleoptericin A (ColA) antimicrobial peptide (AMP) is an important molecular determinant for the maintenance of *S. oryzae/S. pierantonius* homeostasis. By interacting with the bacterial chaperonin GroEL, ColA inhibits bacterial cell septation and generates elongated bacteria with multiple genome copies [18]. Inhibition of *colA* with RNA interference leads to bacterial escape from the bacteriome, and colonization of host surrounding tissues [18]. ColA expression in the bacteriome is dependent on *relish* and *imd*, two genes belonging to the immune deficiency (IMD) pathway [21]. Recently, the weevil’s peptidoglycan recognition protein LB (PGRP-LB) was also shown to play a central role in host homeostasis. By cleaving the tracheal cytotoxin (TCT), a monomeric form of DAP-type peptidoglycan constantly produced by the endosymbionts within the bacteriome, PGRP-LB prevents the exit of TCT from the bacteriome to the insect’s hemolymph, avoiding a chronic activation of host IMD dependent humoral immunity [19]. Taken together, these results suggest that bacterial compartmentalization in the bacteriome is a key strategy that allows the tolerance of symbiotic bacteria as it avoids the contact between the endosymbionts and the insect’s immune system [22], therefore preventing chronic activation of the host immune IMD pathway against the beneficial microorganisms [23].

Current knowledge of gene expression levels in the larval bacteriome is limited to a couple of AMPs and a few other stress-related insect genes [19–21], and little is known about other insect or bacterial regulatory mechanisms involved in endosymbiont protection from bacteriocyte immune activation. We have previously shown that the bacteriome participates in the immune response against pathogenic bacteria and TCT challenge. Notably, up-regulation of several AMPs in weevils after injection of bacteria into the insect hemolymph is observed in the bacteriome [17,20], as well as in the rest of the body [17,20,24]. In addition, TCT injection is sufficient to mimic AMP induction in larval bacteriomes upon bacterial challenge [19]. It is important to note that AMP induction upon TCT challenge is IMD dependent, as is the control of endosymbionts within bacteriocytes, indicating the same pathway can fight exogenous bacterial infection while controlling intracellular beneficial bacteria [21]. Although the involvement of the bacteriome in the immune response would appear in disagreement with its primary function of hosting bacteria, such activation of the immune response against external infections does not seem to pose a threat to *S. pierantonius* integrity since bacterial infections do not induce a reduction in the number of symbionts [20]. This suggests that, despite activating the same immune pathway, differences must exist between fighting external infections and protecting the intracellular symbiont. We hypothesize that either the endosymbionts have evolved specific mechanisms to counteract the bacteriome immune response, or that host-controlled mechanisms, such as AMP secretion, ensure endosymbiont protection.

In this work, we conducted a global dual transcriptomic analysis of host bacteriomes and bacteria challenged systemically with TCT, in order to mimic an immune response in the absence of a real infectious threat. While confirming the involvement of the bacteriome in the immune response, notably via an AMP induction, immunohistochemical observations showed AMP accumulation only outside of the bacteriome, and a full preservation of the basal bacterial transcriptional program. Thus, efficient physical separation between symbionts and bacteria-harnessing molecules ensures full symbiont protection during an immune challenge.

## Methods

### Animal rearing, peptidoglycan challenge, and sample preparation

*S. oryzae* laboratory strain (Bouriz) were reared on wheat grains at 27.5°C and at 70% relative humidity. A strain of aposymbiotic insects was obtained as previously described [25]. The DAP-type peptidoglycan fragment TCT was purified from *Escherichia coli* as previously described [26]. Fourth instar larvae were extracted from wheat grains and challenged with a 0.2 mM TCT solution diluted in 1X phosphate buffered saline (PBS) injected into the hemolymph using a Nanoject III (Drummond). Sterile phosphate buffered saline PBS was also used as a negative control. Injected and non-injected larvae (naïve) were kept in white flour for 6 hours at 27.5°C and at 70% relative humidity before dissection. Bacteriomes were dissected in diethylpyrocarbonate-treated Buffer A (25 mM KCl, 10 mM MgCl_2_, 250 mM sucrose, 35 mM Tris/HCl, pH = 7.5). For each sample, bacteriomes were pooled (30 for Dual RNA-seq library preparation, and at least 25 for RT-qPCR), and stored at −80 °C prior to RNA extraction. Pools of five carcasses from symbiotic dissected weevils were used for RT-qPCR. Aposymbiotic samples consisted in pools of five fourth instar aposymbiotic larvae, which were torn in Buffer A, but not dissected as they do not harbor bacteriomes.

### RNA extraction, library preparation and sequencing

Total RNA was extracted with TRIzol™ Reagent (Invitrogen, ref.: 15596026) following the manufacturer’s instructions. Nucleic acids were then purified using the NucleoSpin RNA Clean up kit (Macherey Nagel, ref.: 740948). Genomic DNA was removed from the samples with the DNA free DNA removal kit (Ambion, ref.: AM1906). Total RNA concentration and quality were checked using the Qubit Fluorometer (ThermoFisher Scientific) and Tapestation 2200 (Agilent Biotechnologies). Ribo-depletion and Dual RNA-seq strand-specific cDNA libraries were obtained starting from 100 ng of total RNA using the Ovation Universal RNA-seq System (NuGEN) following the manufacturer’s instructions. Libraries were sequenced on a Nextseq 500 sequencer (Illumina), using the NextSeq 500/550 High Output Kit (Illumina).

### Preprocessing, mapping of reads and differential expression analysis

Raw reads were processed using Cutadapt v1.18 [27] to remove adapters, filter out reads shorter than 50 bp and reads that had a mean quality value lower or equal to 30. Clean reads were mapped against the *S. oryzae* genome (Genbank: PRJNA431034) with STAR v2.7.3a [28], and against the S*. pierantonius* genome (Genbank: CP006568.1) with Bowtie 2 v2.3.5 [29] with default parameters. Shared reads between the two genomes were filtered out with the aid of SAMtools v1.10 [30] and Picard v2.21.6 (available from https://broadinstitute.github.io/picard/). Gene counts were obtained for uniquely mapped reads with featureCounts v1.6.4 method from the Subread package [31]. Whenever uniquely mapped read counts were set to zero due to duplicated regions or multi-mapped reads, we further verified these regions within the multi-mapped read counts available with featureCounts. Insertion sequence (IS) families from the bacteria were also counted with the use of TEtools (v1.0.0) with default parameters [32]. Gene counts and TEtools counts were used as input for differential expression analyses using the DESeq2 v1.26.0 [33] package in R. After testing, the p-values were adjusted with the Benjamini-Hochberg correction [34] for multi-testing. Genes were considered differentially expressed when adjusted p-values (p-adj) were smaller than 0.05. Sequencing data from this study have been deposited at the National Center for Biotechnology Information Sequence Read Archive, https://www.ncbi.nlm.nih.gov/sra (accession no. PRJNA816415).

### Quantitative RT-PCR

Total RNA was extracted from fourth instar bacteriomes and carcasses, as well as from whole aposymbiotic fourth instar larvae using the RNAqueous - Micro kit (Ambion). DNA was removed with DNAse treatment and RNA quality was checked with Nanodrop (ThermoFisher Scientific). Complementary DNA (cDNA) was produced with the iScript™ cDNA Synthesis Kit (Bio-Rad) following the manufacturer’s instructions and starting with 500 ng total RNA. Differential gene expression was assessed by quantitative real-time PCR with a CFX Connect Real-Time PCR Detection System (Bio-Rad) using the LightCycler Fast Start DNA Master SYBR Green I kit (Roche Diagnostics), as previously described [19], except for *dpt4*, for which the annealing temperature was reduced to 54.5 °C. Data were normalized using the ratio of the target cDNA concentration to the geometric average of two housekeeping transcripts: *glyceraldehyde 3-phosphate dehydrogenase* (LOC115881082) and *malate oxidase* (LOC115886866). Primers were designed to amplify fragments of approximately 150 bp. A complete list of primers can be found in Additional Table 1.

### Immunohistochemi stry

Larval samples challenged with TCT or PBS were prepared for histological observations as described in [19]. Briefly, samples were fixed in paraformaldehyde (PFA) 4%. After one day, the fixative was replaced by several washings with PBS before embedding the tissue in 1.3% agar, then dehydrated through a gradient of ethanol (EtOH) washes and transferred to butanol-1, at 4°C, overnight. Samples were then placed in melted Paraplast and 3 μm-thick sections were cut with a HM 340 E microtome (ThermoFisher Scientific). Sections were placed on poly-lysine-coated slides, dried overnight at 37°C, and stored at 4°C.

For AMP localization, samples were dewaxed twice in methylcyclohexane for 10 min, rinsed in EtOH 100°, rehydrated through an EtOH gradient and then placed in PBS with 1% Bovine Serum Albumin (BSA) for 30 min. ColA rabbit primary polyclonal anti-serum (Login et al., 2011) at 1:200 dilution, and a Coleoptericin B (ColB) primary polyclonal anti-serum (Proteogenix, Schiltigheim-France) at 1:300 dilution in 0.1% BSA were used. Preimmune rabbit serum (J0) was used as a negative control for ColA anti-serum, and BSA 0.1% for ColB (purified antibody). Antibody specificity was checked by western blot. After 1 h incubation at room temperature in the dark, sections were washed with PBS containing 0.2% Tween. Samples were then incubated with anti-rabbit IgG, labeled with Alexa Fluor 488. This secondary antibody was applied for 1 h at room temperature, diluted at 1:500 in 0.1% BSA in PBS. The excess of secondary antibody was washed with PBS-Tween, rinsed with PBS and washed several times with tap water. Sections were then dried and mounted using PermaFluorTM Aqueous Mounting Medium (ThermoFisher Scientific), together with 4,6-diamidino-2-phenylindole (DAPI, Sigma-Aldrich) for nuclear staining (3 μg/ml of medium). Images were acquired using an epifluorescence microscope (Olympus IX81), under specific emission filters: HQ535/50 for the green signal (antibody staining), D470/40 for the blue signal (DAPI) and HQ610/75 for the red signal (unspecific autofluorescence from tissue). Images were captured using an XM10 camera and the CellSens Software (Soft Imaging System). Images were treated using ImageJ (release 1.47v).

## Results and discussion

### Dual RNA sequencing successfully yielded both insect and bacterial transcripts

To investigate bacteriome response to an immune challenge, we extracted *S. oryzae*’s fourth instar larvae (L4) from grains and injected them with TCT, a fragment of the DAP-type peptidoglycan produced by Gram-negative bacteria, including *S. pierantonius* [19] and recovered bacteriomes six hours post injection as previously described [20]. TCT injection is able to trigger a potent response without the interference of an exogenous infectious bacteria [21]. Control larvae were injected with PBS, or extracted from grains but not injected (See Figure 1). To obtain the transcriptomic profile of both the symbiont and the host, Dual RNA-seq was performed in triplicates and yielded from 105 to 140 M reads per library (Additional Table 2). The reads were cleaned from adapter sequences and low-quality reads, and around 85% of the raw reads were kept for further analyses. We subsequently mapped the clean reads against both genomes, and obtained ~65-80% unambiguously mapping to the genome of *S. oryzae*, and ~5-8% to the genome of *S. pierantonius*. In each library from 23 to 33 M reads were uniquely mapped against insect genes (Additional Table 3), whereas ~3 M reads were uniquely mapped against bacterial genes (Additional Table 4). These results depict an improvement from our previous study, which yielded ~0.4 M reads mapped against bacterial genes in the same developmental stage and similar sequencing depth [16].

**Figure 1.**
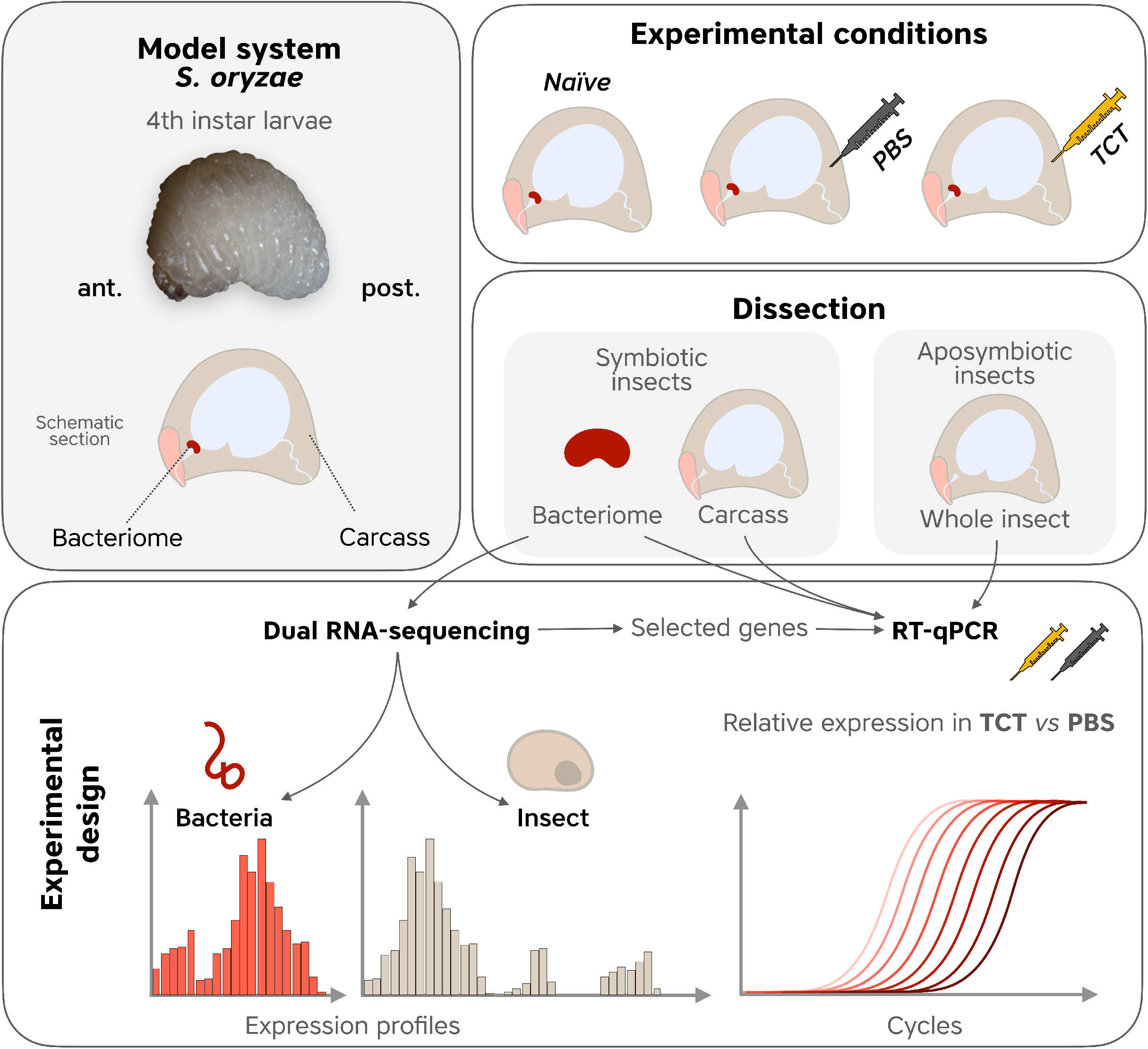
Schematic diagram of the experimental design. Top left panel: image of a *S. oryzae* 4th instar larva along with a schematic section. Top left panels: symbiotic and aposymbiotic *S. oryzae* fourth instar larvae were extracted from grains and dorsally injected with 0.2 mM PBS, 0.2 mM TCT at the level of the haemolymph. Other larvae were extracted from grains but not injected (Naïve). Bacteriomes and carcasses were sampled from PBS/TCT-injected or naïve symbiotic larvae alongside whole aposymbiotic larvae. Bottom panel: dual RNA-seq was performed to detect insect and bacterial expression profiles was performed on bacteriomes and carcasses of symbiotic weevils (PBS, TCT and Naïve samples). RT-qPCR experiments were performed on TCT and PBS-treated bacteriomes/carcasses from symbiotic weevils as well as whole larvae from aposymbiotic weevils, to detect bacteriome-specific and/or symbiont-dependent transcriptomic changes.

### Systemic TCT challenge triggers AMP induction within the bacteriome

Sixteen *S. oryzae* genes were detected as differentially expressed (DE; p-adj < 0<.05) six hours after the TCT challenge in the bacteriome, with respect to the bacteriome of non-injected (naïve) or PBS-injected larvae (Table 1, Additional Table 5). Among these, one gene was strongly down-regulated, four were mildly down-regulated, and eleven were up-regulated in response to TCT.

**Table 1:** *S. oryzae* genes differentially expressed in TCT-challenged bacteriomes (p-adj < 0.05) identified by Dual RNA-seq.

| Gene Information |  |  |  | Average expression (TPM)* |  |  | Log2 Fold Change |  |
| --- | --- | --- | --- | --- | --- | --- | --- | --- |
| Gene ID | Type | Gene Abbreviation | Protein Name | Naïve | PBS | TCT | TCT vs Naïve | TCT vs PBS |
| *LOC115882681* | Unknown | N/A | Uncharacterized | 2.93 | 1.06 | 0.03 | -6.941 | -5.574 |
| *LOC115888453* | Transcription Factor | *adf-1* | Transcription factor Adf-1 family | 54.55 | 49.23 | 34.33 | -0.696 | -0.601 |
| *LOC115881033* | Translation Initiation | *eif4ebp-2* | Eukaryotic translation initiation factor 4E-binding protein 2 | 256.09 | 249.36 | 162.63 | -0.691 | -0.706 |
| *LOC115891903* | Transcription Factor | *nrbp* | Nuclear receptor-binding protein | 117.12 | 128.47 | 83.03 | -0.518 | -0.600 |
| *LOC115883362* | Transcription Factor | *znf-91* | Zinc finger protein 91-like | 48.09 | 46.24 | 36.16 | -0.436 | -0.434 |
| *LOC115877563* | ABC Transporter | *mrp-4* | Multidrug resistance-associated protein 4-like | 1.41 | 1.30 | 3.44 | 1.277 | 1.335 |
| *LOC115885681* | Growth Factor | *brx* | Barietin toxin | 1.24 | 1.13 | 4.30 | 1.793 | 1.881 |
| *LOC115886735* | Bacterial Recognition | *gnbp-1* | Beta-1,3-glucan-binding protein-like | 7.07 | 8.72 | 37.97 | 2.411 | 2.052 |
| *LOC115874620* | AMP | *col-A* | Coleopteracin-A | 251.44 | 480.30 | 2116.35 | 3.030 | 2.040 |
| *LOC115883884* | AMP | *lux* | Luxuriosin | 7.34 | 6.01 | 60.94 | 3.042 | 3.256 |
| *LOC115884866* | AMP | *glyr-amp* | Glycine-Rich AMP | 11.10 | 12.88 | 120.87 | 3.410 | 3.165 |
| *LOC115888387* | AMP | *srx* | Sarcotoxin | 27.81 | 28.61 | 407.20 | 3.826 | 3.734<sup>s</sup> |
| *LOC115877462* | AMP | *dpt-2* | Diptericin-2 | 40.45 | 63.50 | 731.07 | 4.131 | 3.425 |
| *LOC115877463* | AMP | *dpt-3* | Diptericin-3 | 9.45 | 13.71 | 261.58 | 4.759 | 4.164 |
| *LOC115874703* | AMP | *col-B* | Coleopteracin-B | 1.97 | 3.42 | 86.46 | 5.386 | 4.538 |
| *LOC115877465* | AMP | *dpt-4* | Diptericin-4 | 1.31 | 2.52 | 98.43 | 6.196 | 5.225 |

§: This transcript was below the significance of detection in one of the conditions due to an outlier; results were verified with EdgeR and we validated this as a DE gene after the qPCRs.
\* Average expression is provided in transcripts per million (TPM).

RT-qPCR experiments confirmed the TCT-dependent induction of all 11 up-regulated genes (Figure 2, Additional Figure 1). Eight of these genes encode AMPs and all possess a predicted signal peptide: *colA* (Coleoptericin A), Coleoptericin B (*colB*), Sarcotoxin (*srx*), Luxoriosin (*lux*), a Gly-rich AMP (*gly-rich AMP*), and three Diptericins (*dpt-2, 3* and *4*, Figure 2) [35]. This AMP induction is in agreement with previous reports, where AMPs induced in larvae by immune challenge included *colA* [17,20,21,24], *colB*, *srx* [20,21,24], *dpt*, *cecropin* and *defensins* [20,24]. In addition to the eight AMPs, genes encoding one Gram-negative binding protein (*gnbp-2*), a barietin-like toxin (*brx*) and a multidrug resistant protein (*mrp-4*) were also up-regulated in the bacteriome (Figure 3). These three genes have not been identified in previous studies. *gnbp-2* is likely involved in insect defense responses against Gram-negative bacteria [36,37] and, like AMPs, contains a predicted secretory sequence at the peptide N-terminus (SignalP 6.0 likelihood value of 0.9998). It is noteworthy that another member of the *gnbp-2* family was also shown to be up-regulated in *S. oryzae* bacteriome in response to a bacterial challenge in a previous study [24]. The barietin-like toxin likely acts as a toxin directed against bacteria [38], similarly to AMPs, and also contains a predicted secretory sequence in the N-terminal region (SignalP 6.0 likelihood value of 1.0). Finally, *mrp-4 like* is likely a transporter involved in secretion of toxin and/or regulating homeostasis against pathogens [39]. In contrast, none of the down-regulated bacteriome genes detected in the Dual RNA-seq were confirmed by RT-qPCR (Additional Figure 1). These results might be explained by their less pronounced down-regulation as seen by a milder Log2FC. Moreover, Dual RNA-seq was obtained from total ribodepleted RNAs, while RT-qPCR was performed on polyadenylated mRNAs, which could contribute to the differences observed in these analyses.

**Figure 2.**
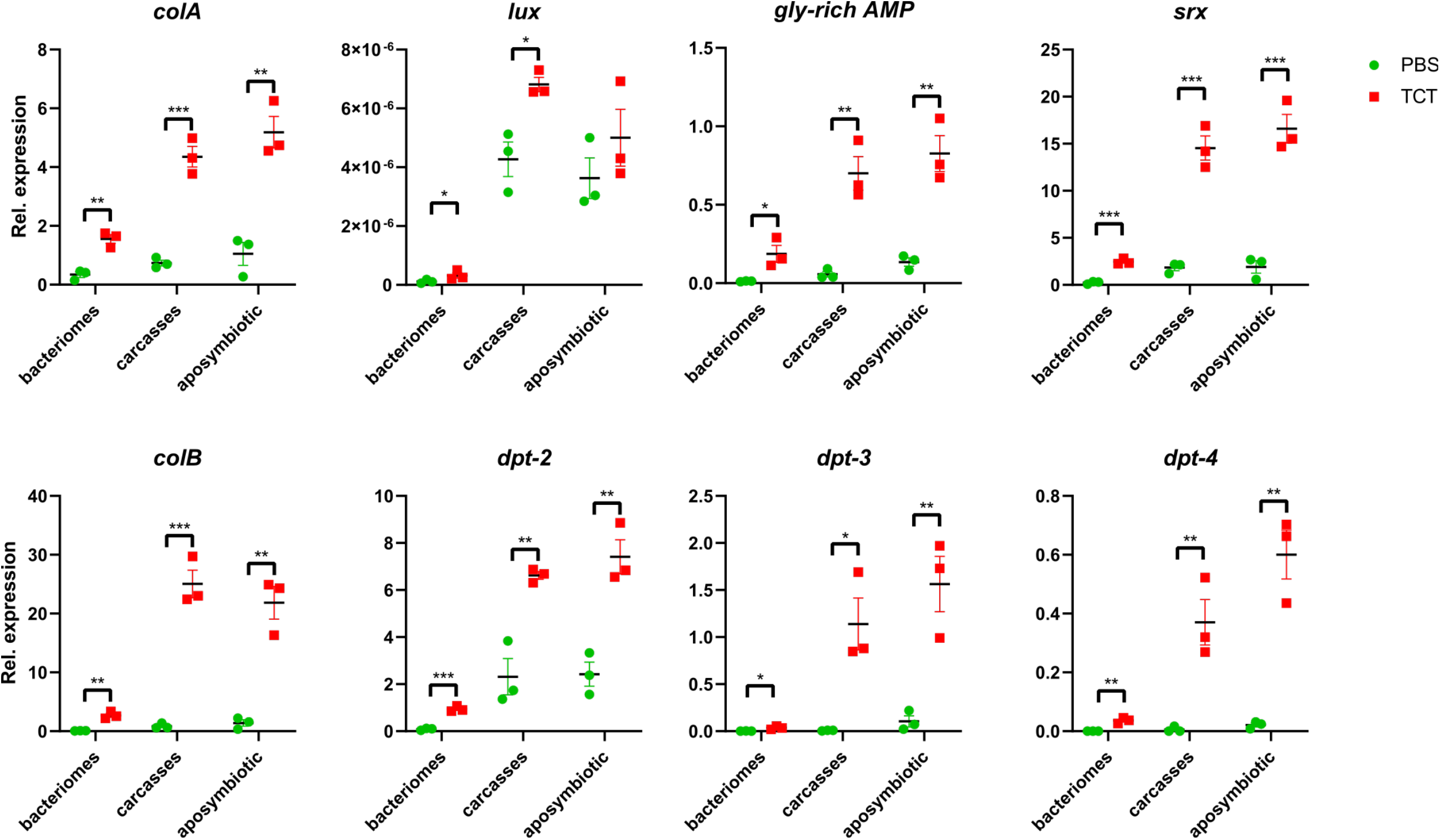
Differential expression of TCT-induced AMPs in bacteriomes. The quantification was performed by RT-qPCR on *S. oryzae* bacteriomes and carcasses of symbiotic weevils, as well as on whole aposymbiotic larvae. Green dots: PBS-injected larvae (control); red squares: TCT-injected larvae. Asterisks denote statistical significance (ANOVA with Kruskal-Wallis test, * = p ≤ 0.05). Error bars represent SE. Overall, the AMP induction in response to TCT is observed in both bacteriomes and carcasses of symbiotic weevils, as well as in aposymbiotic weevils.

**Figure 3.**
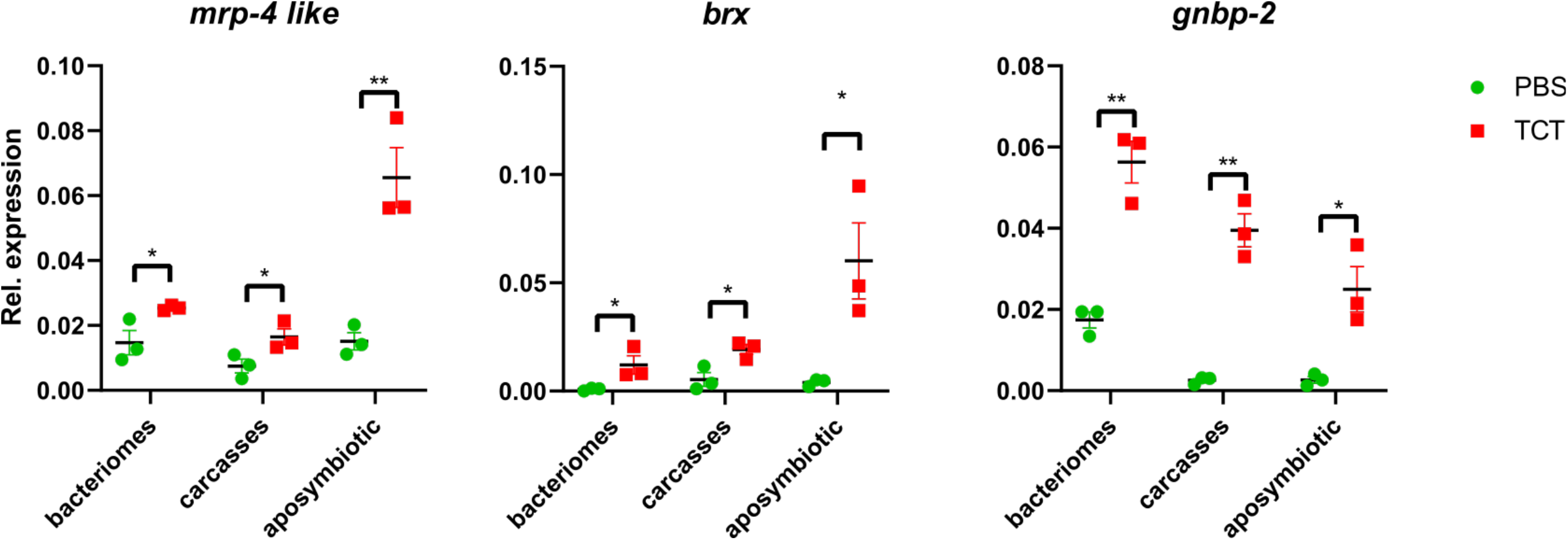
Differential expression of TCT-induced genes in bacteriomes other then AMPs. The quantification was performed by RT-qPCR on *S. oryzae* bacteriomes and carcasses of symbiotic weevils, as well as on whole aposymbiotic larvae. Green dots: PBS-injected larvae (control); red squares: TCT-injected larvae. Asterisks denote statistical significance (ANOVA with Kruskal-Wallis test, * = p ≤ 0.05). Error bars represent SE. Overall, upregulation in response to TCT is observed in both bacteriomes and carcasses of symbiotic weevils, as well as in aposymbiotic weevils.

To test whether the identified up-regulated genes were part of a bacteriome-specific response, we analyzed the expression of the same genes in TCT- or PBS-challenged carcasses of symbiotic insects as well as in TCT- or PBS-challenged aposymbiotic L4 (*i.e.* insects artificially devoid of symbionts, with no bacteriome, see *Methods* Section). We found that all eight up-regulated AMPs (Figure 2) and the other three up-regulated genes (Figure 3) were also induced in TCT-challenged symbiotic carcasses, and TCT-challenged aposymbiotic whole larvae. The steady-state gene levels in PBS injection were comparable between the three conditions, with the exception of *lux*, *dpt-3* and *srx*. Finally, in agreement with previous studies, these data show that the bacteriome induction is generally milder than the systemic response [20], but confirms the involvement of the bacteriome in the host immune response. Previous studies have shown that *colA* is chronically expressed in the larval bacteriomes, here seen at ~250 transcripts per million (TPM) in control conditions, and it successfully prevents endosymbiont escape and morphology (Login et a., 2011, Maire et al., 2019). The TCT-induced AMP upregulation in the bacteriomes might therefore constitute a threat for endosymbiont fitness. Overall, these results strongly suggest that the presence of *S. pierantonius* does not affect the systemic induction of AMPs, which is comparable between symbiotic and aposymbiotic insects.

It is important to note that the present study failed to detect a couple of host genes previously identified as up-regulated upon bacterial infection in *S. oryzae*, including the regulatory gene *pirk*, and the Toll pathway-related genes (*pgrp*, *toll*), among others [20]. These discrepancies might indicate the inability of the TCT molecule to trigger a complete immune response, as opposed to a whole bacterium. TCT is a monomeric form of DAP-type PG and induces only the IMD and not the Toll pathway [19]. Nevertheless, the AMP induction observed here is consistent with previous studies [20,24] and would be expected to constitute a severe threat for the endosymbionts in the absence of protective mechanisms.

### Symbiotic bacteria are insensitive to the activation of the bacteriome immune system

In order to identify potential signatures of bacterial stress and gene modulations to counteract the insect immune response and AMP induction, the symbiont transcriptomic profile obtained by Dual RNA-seq from TCT-challenged bacteriome samples was compared with controls, *i.e.* PBS-injected or naïve. Remarkably, the differential analysis revealed that bacterial transcription is unresponsive to the TCT challenge (Additional Tables 6). Furthermore, and similarly to coding regions, we did not detect changes in expression in repetitive regions (IS) (Additional Table 7). Moreover, a previous study using Dual RNA-seq in *S. oryzae* showed around 400 differentially expressed bacterial genes throughout the metamorphosis of the insect, confirming the ability of the endosymbiont to modulate gene expression in response to host developmental stimuli [16]. The contrast between large changes of gene expression during metamorphosis, with a complete lack of differentially expressed genes upon TCT challenge, strongly suggests that the bacteria do not sense the AMP induction or any other stress induced by such challenge [42].

Rather, analysis of the complete bacterial transcriptome from both controls and TCT-challenged larvae display similar gene expression. Highly expressed bacterial protein coding genes detected within the bacteriome (Additional Table 8) are mainly involved in transcriptional regulation, translation, stress response, and virulence (see Additional Text 1 for more information). Several transcriptional, translational and stabilization factors of the general stress response sigma factor RpoS (reviewed in [40]) were similarly expressed at varied levels in all conditions (Additional Table 9). The expression of *rpoS* was lower than the vegetative sigma factor *rpoD*, which is a typical profile of the exponential growth phase in *Escherichia coli* [41]. This basal level of *rpoS* is also needed for triggering a fast stress response in diverse bacteria [40], and shows the ability of *S. pierantonius* from larval bacteriomes to quickly enter a “virulent mode” in the subsequent pupal stage that allows them to exit bacteriocytes and re-infect stem cells [16].

Together with previous findings that the symbiont population remains unchanged even after an immune challenge with pathogenic bacteria [20], this suggests that other regulatory mechanisms are in place to maintain the physical integrity of the symbiotic bacterial population during host AMP induction.

### Mature AMPs are physically separated from endosymbionts

One of the hallmarks of AMPs is the presence of a N-terminal secretory sequence that addresses them to the outside of the cell, including the hemolymph, to counteract systemic infections [52]. Thus, even though cells in the bacteriome can produce AMPs, their final localization outside of bacteriocytes would ensure protection of the endosymbionts from AMP harm. However, in physiological conditions, ColA is produced by and retained inside the bacteriocytes, together with the endosymbionts, where it keeps them from escaping [18]. Since our knowledge of AMP localization is still limited because of the lack of specific antibodies, it cannot be excluded that other AMPs might also accumulate intracellularly, especially if highly expressed, and constitute a threat for the endosymbionts. We therefore assessed the localization of TCT-induced AMPs with respect to the symbionts. We performed immunohistochemistry with polyclonal antibodies able to recognize *colB*, an AMP previously shown to be induced by TCT and bacterial challenges [19] and the bacteriome-specific AMP ColA [18]. The choice of *colB*, in particular, was dictated by the fact that, despite this peptide being very similar to *colA* (46.72% of amino acid sequence identity), their function is remarkably different, as *colA* is expressed constitutively in the bacteriome where it interacts with GroEL and contributes to the insect-bacteria homeostasis. Samples were taken at six hours after the immune challenge with TCT or PBS (as for the transcriptomic analysis), so that we could confirm that AMPs were induced at the protein level, despite the lack of endosymbiont response. In the PBS-injected controls (Figure 4A-D), ColA was detected within the bacteriome (Figure 4A) - as expected because of its role in preventing symbiont escape (Login et al., 2011) - but not in the other tissues (Figure 4B). These results confirm the presence of ColA within the bacteriome, even at basal expression levels. In contrast, ColB was not detected inside the bacteriome (Figure 4C), nor in other surrounding cells, including gut tissues (Figure 4D). In response to TCT (Figure 4E-H), ColA was still clearly detectable within the bacteriome (Figure 4E), as expected, but also in several epithelial gut cells as well as in the acellular extended region that likely corresponds to the hemolymph (Figure 4F). This confirms the dual role of ColA in both symbiosis control [18] and in response to an exogenous immune challenge. On the contrary, ColB was still absent from the bacteriome tissue following TCT challenge (Figure 4G), but, similarly to ColA, was detected in the hemolymph of TCT-challenged larvae (Figure 4H).

**Figure 4.**
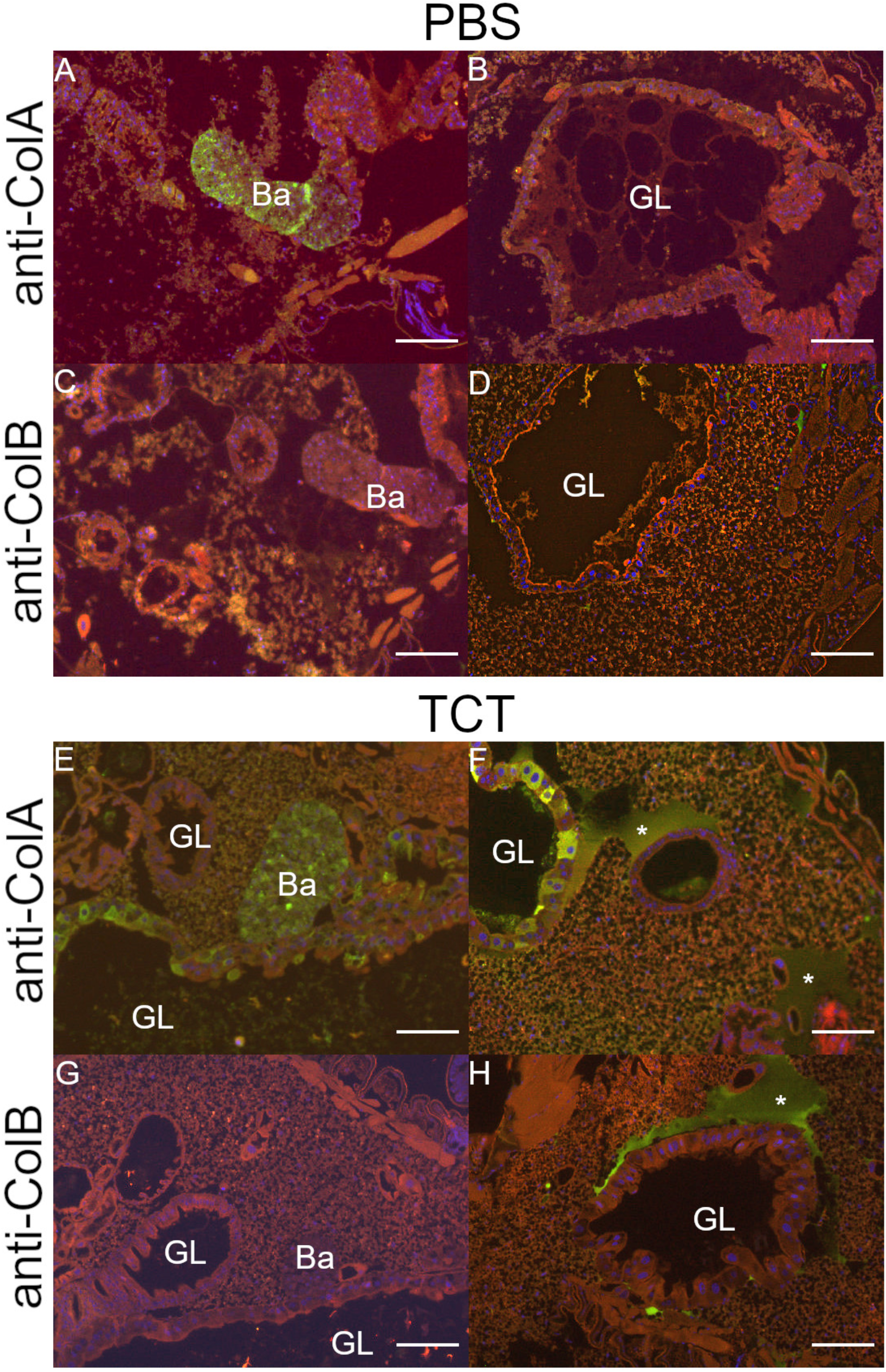
AMP localization in *S. oryzae* larvae, before and after TCT immune challenge. Upper panel: ColA (first row) and ColB (second row) localization in PBS-injected larvae. Lower panel: ColA (first row) and ColB (second row) localization in TCT-injected larvae. Ba: bacteriome; GL: gut lumen. Asterisks indicate accumulation of AMPs in the hemolymph. Scale bar: 50 μm.

The results show that, in agreement with the lack of endosymbiont transcriptomic response, the excess of ColA, ColB (and potentially all other AMPs (whether induced in the bacteriome or the fat body), remain physically separated from the endosymbionts. Thus endosymbiont integrity is protected even while AMPs are participating in the systemic immune response.

## Conclusions

There are currently three main known strategies allowing symbiotic microorganisms to coexist with efficient and responsive insect immunity: *i)* evolution of the ability to differentiate between pathogenic and symbiotic MAMPs by the host, ii) bacterial molecular modifications leading to immune tolerance, notably promoting biofilm formation [53], and *iii)* compartmentalization of the symbionts in specialized symbiotic organs, often called bacteriomes [54]. The compartmentalization strategy sequesters the symbionts in specialized cells, creating a favorable environment for their metabolic activity, and keeping them under control while avoiding overproliferation and virulence. The bacteriomes are therefore found in many insect species, including aphids [55], planthoppers [56], cicadas [57], and beetles [58]. Although very common, little is known about the evolution and immune modulation inside the bacteriomes, as well as on their formation and maintenance.

In the *S. oryzae*/*S. pierantonius* symbiosis, bacterial MAMPs are able to trigger a potent immune response, thus excluding a selective tolerance of the weevil immune system towards *S. pierantonius* MAMPs [17,19–21]. The absence of bacterial transcriptomic response to the systemic TCT immune challenge excludes active mechanisms of immune suppression from the endosymbiont. Rather, compartmentalization of *S. pierantonius* within bacteriomes guarantees physical separation between the endosymbionts and AMPs that might be produced by the bacteriome itself or elsewhere (*e.g.* fat bodies). This mechanism is crucial to protect both the host from the symbionts, and the bacteria from the insect immune system [18,21]. As demonstrated by the immunofluorescence labeling, there is no colocalization of endosymbiont-containing cells and AMPs, with the notable exception of ColA due to its homeostatic function, thus showing that not only the bacteriome acts as a physical barrier against the external AMPs, but is also capable to efficiently drain away the toxic molecules produced both inside or outside the bacteriome (Figure 5). Altogether these data refine the understanding on how an organ such as the bacteriome can ensure specific symbiotic function, *i.e.* maintain and control endosymbionts in a specific location, while potentially participating in the immune response to exogenous bacteria.

**Figure 5.**
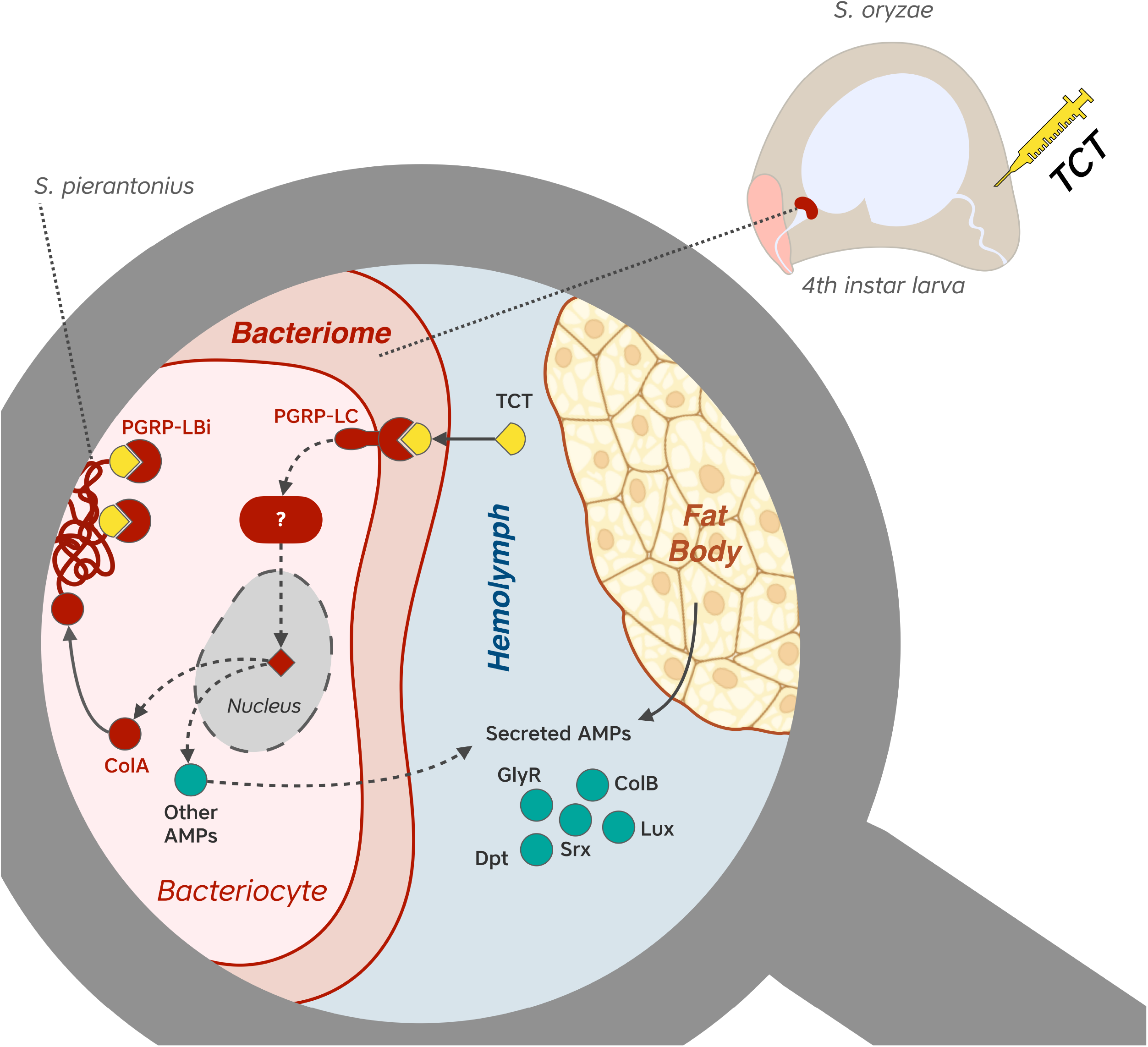
Proposed mechanisms of TCT challenge response within bacteriomes of *S. oryzae.* TCT injected in the hemolymph reaches bacteriomes and is recognized by PGRP-LC from bacteriocytes. Through a signaling cascade potentially dependent on IMD/RELISH proteins, bacteriocytes activate an AMP induction which for the most part are thought to be secreted (ColB, Srx, Lux, Gly-Rich AMP, Dpt-2, −3, and 4) to aid in the global immunity response, but no effectors are perceived by the bacteria within bacteriomes. AMPs are also produced by the fat body and secreted to the hemolymph. ColA in turn is kept within bacteriocytes to prevent *S. pierantonius* from exiting the host cells during this immune challenge.

## List of abbreviations

AMP: (antimicrobial peptide)
BSA: (bovine serum albumin)
cDNA: (complementary DNA)
DAPI: (4,6-diamidino-2-phenylindole)
EtOH: (ethanol)
IMD: (immune deficiency)
IS: (insertion sequence)
L4: (fourth instar larvae)
MAMPs: (microbial-associated molecular patterns)
p-adj: (adjusted p-values)
PBS: (phosphate buffered saline)
PFA: (paraformaldehyde)
PG: (proteoglycan)
T3SS: (type III secretion system)
TA: (toxinantitoxin)
TCT: (tracheal cytotoxin)
TPM: (transcripts per million)

## Declarations

### Ethical approval and consent to participate

Not applicable

### Consent for publication

Not applicable

### Availability of data and materials

Sequencing data from this study have been deposited at the National Center for Biotechnology Information Sequence Read Archive, https://www.ncbi.nlm.nih.gov/sra (accession no. PRJNA816415).

### Author’s contributions

AH, CV and RR conceived the original project. EDA was responsible for all molecular biology methods, with the help of AV. AV in collaboration with SH and BG constructed the Dual RNA-seq libraries and produced the sequencing reads. MGF was responsible for the bioinformatic analyses of the Dual RNA-seq with the help of NP. EDA, MGF and RR analyzed the data. EDA with the help of SB performed the immunofluorescence experiments. EDA, MGF, RR wrote the manuscript with the help of CV, AH, CVM and AZR. All authors read and approved the final manuscript.

## Acknowledgements

We are grateful to Dr. Dominique Mengin-Lecreulx for providing us with purified TCT and TCT preparation protocol. We are also grateful to Dr. Julien Orlans who subsequently took charge of TCT production. We thank Marie-France Sagot for the insightful discussions regarding the dual transcriptomic analyses and two anonymous reviewers for their constructive feedback.

## Author’s information

Twitter handles: @AnnaZaidmanRemy (Anna Zaidman-Rémy), @cmonegat (Carole Vincent-Monegat), @Cosmicomica (Elisa Dell’Aglio), @MGFerrarini (Mariana Galvão Ferrarini), @niparisot (Nicolas Parisot) and @rita_rebollo (Rita Rebollo).

## Additional Files

Additional Figure 1. Differential expression of TCT-repressed genes in bacteriomes, according to Dual RNA-seq. The quantification was performed by qRT-PCR on *S. oryzae* bacteriomes and carcasses of symbiotic weevils, as well as on whole aposymbiotic larvae. Green dots: PBS-injected larvae (control); red squares: TCT-injected larvae.

Additional Text 1: Description of genes highly expressed *from S. pierantonius*.

Additional Table 1: Primer sequences

Additional Table 2: Dual RNA-seq trimming and mapping statistics.

Additional Table 3: Count data of *S. oryzae* genes

Additional Table 4: Differential expression analysis of *S. oryzae* genes.

Additional Table 5: Level of expression of *S. pierantonius* genes.

Additional Table 6: Differential expression of *S. pierantonius* genes.

Additional Table 7: Count data and differential expression analysis of *S. pierantonius* ISs.

Additional Table 8: Highly expressed bacterial genes in all conditions (TPM > 1000) belonging to key biological functions in *S. pierantonius*.

Additional Table 9: Expression levels of genes related to the general stress response in S. pierantonius.

## References

1. Moran NA. Symbiosis. Curr Biol CB. 2006;16: R866–871. doi:10.1016/j.cub.2006.09.019

2. Moya A, Peretó J, Gil R, Latorre A. Learning how to live together: genomic insights into prokaryote–animal symbioses. Nat Rev Genet. 2008;9: 218–229. doi:10.1038/nrg2319

3. Heddi A, Grenier A-M, Khatchadourian C, Charles H, Nardon P. Four intracellular genomes direct weevil biology: Nuclear, mitochondrial, principal endosymbiont, and Wolbachia. Proc Natl Acad Sci. 1999;96: 6814–6819. doi:10.1073/pnas.96.12.6814

4. Tsuchida T, Koga R, Fukatsu T. Host Plant Specialization Governed by Facultative Symbiont. Science. 2004;303: 1989–1989. doi:10.1126/science.1094611

5. Wilson ACC, Ashton PD, Calevro F, Charles H, Colella S, Febvay G, et al. Genomic insight into the amino acid relations of the pea aphid, Acyrthosiphon pisum, with its symbiotic bacterium Buchnera aphidicola. Insect Mol Biol. 2010;19: 249–258. doi:10.1111/j.1365-2583.2009.00942.x

6. Aksoy S, Caccone A, Galvani AP, Okedi LM. Glossina fuscipes populations provide insights for human African trypanosomiasis transmission in Uganda. Trends Parasitol. 2013;29: 394–406. doi:10.1016/j.pt.2013.06.005

7. Zaidman-Rémy A, Vigneron A, Weiss BL, Heddi A. What can a weevil teach a fly, and reciprocally? Interaction of host immune systems with endosymbionts in Glossina and Sitophilus. BMC Microbiol. 2018;18: 150. doi:10.1186/s12866-018-1278-5

8. Zug R, Hammerstein P. Wolbachia and the insect immune system: what reactive oxygen species can tell us about the mechanisms of Wolbachia–host interactions. Front Microbiol. 2015;6. doi:10.3389/fmicb.2015.01201

9. He Z, Wang P, Shi H, Si F, Hao Y, Chen B. Fas-associated factor 1 plays a negative regulatory role in the antibacterial immunity of Locusta migratoria. Insect Mol Biol. 2013;22: 389–398. doi:10.1111/imb.12029

10. Lefèvre C, Charles H, Vallier A, Delobel B, Farrell B, Heddi A. Endosymbiont phylogenesis in the dryophthoridae weevils: evidence for bacterial replacement. Mol Biol Evol. 2004;21: 965–973. doi:10.1093/molbev/msh063

11. Clayton AL, Oakeson KF, Gutin M, Pontes A, Dunn DM, Niederhausern AC von, et al. A Novel Human-Infection-Derived Bacterium Provides Insights into the Evolutionary Origins of Mutualistic Insect– Bacterial Symbioses. PLOS Genet. 2012;8: e1002990. doi:10.1371/journal.pgen.1002990

12. Mansour K. Memoirs: Preliminary Studies on the Bacterial Cell-mass (Accessory Cell-mass) of Calandra Oryzae (Linn.): The Rice Weevil. J Cell Sci. 1930;s2-73: 421–435. doi:10.1242/jcs.s2-73.291.421

13. Vigneron A, Masson F, Vallier A, Balmand S, Rey M, Vincent-Monégat C, et al. Insects Recycle Endosymbionts when the Benefit Is Over. Curr Biol. 2014;24: 2267–2273. doi:10.1016/j.cub.2014.07.065

14. Grenier AM, Nardon C, Nardon P. The role of symbiotes in flight activity of Sitophilus weevils. Entomol Exp Appl. 1994;70: 201–208. doi:10.1111/j.1570-7458.1994.tb00748.x

15. Oakeson KF, Gil R, Clayton AL, Dunn DM, von Niederhausern AC, Hamil C, et al. Genome Degeneration and Adaptation in a Nascent Stage of Symbiosis. Genome Biol Evol. 2014;6: 76–93. doi:10.1093/gbe/evt210

16. Maire J, Parisot N, Galvao Ferrarini M, Vallier A, Gillet B, Hughes S, et al. Spatial and morphological reorganization of endosymbiosis during metamorphosis accommodates adult metabolic requirements in a weevil. Proc Natl Acad Sci. 2020;117: 19347–19358.

17. Anselme C, Pérez-Brocal V, Vallier A, Vincent-Monegat C, Charif D, Latorre A, et al. Identification of the Weevil immune genes and their expression in the bacteriome tissue. BMC Biol. 2008;6: 43. doi:10.1186/1741-7007-6-43

18. Login FH, Balmand S, Vallier A, Vincent-Monégat C, Vigneron A, Weiss-Gayet M, et al. Antimicrobial Peptides Keep Insect Endosymbionts Under Control. Science. 2011;334: 362–365. doi:10.1126/science.1209728

19. Maire J, Vincent-Monégat C, Balmand S, Vallier A, Hervé M, Masson F, et al. Weevil pgrp-lb prevents endosymbiont TCT dissemination and chronic host systemic immune activation. Proc Natl Acad Sci. 2019;116: 5623–5632. doi:10.1073/pnas.1821806116

20. Masson F, Vallier A, Vigneron A, Balmand S, Vincent-Monégat C, Zaidman-Rémy A, et al. Systemic Infection Generates a Local-Like Immune Response of the Bacteriome Organ in Insect Symbiosis. J Innate Immun. 2015;7: 290–301. doi:10.1159/000368928

21. Maire J, Vincent-Monégat C, Masson F, Zaidman-Rémy A, Heddi A. An IMD-like pathway mediates both endosymbiont control and host immunity in the cereal weevil Sitophilus spp. Microbiome. 2018;6: 6. doi:10.1186/s40168-017-0397-9

22. Tsakas S, Marmaras VJ. Insect immunity and its signalling: an overview. Invertebr Surviv J. 2010;7: 228–238.

23. Ratzka C, Liang C, Dandekar T, Gross R, Feldhaar H. Immune response of the ant Camponotus floridanus against pathogens and its obligate mutualistic endosymbiont. Insect Biochem Mol Biol. 2011;41: 529–536. doi:10.1016/j.ibmb.2011.03.002

24. Vigneron A, Charif D, Vincent-Monégat C, Vallier A, Gavory F, Wincker P, et al. Host gene response to endosymbiont and pathogen in the cereal weevil Sitophilus oryzae. BMC Microbiol. 2012;12: S14. doi:10.1186/1471-2180-12-S1-S14

25. Nardon P. Obtention d’une souche asymbiotique chez le charançon Sitophilus sasakii Tak: différentes méthodes d’obtention et comparaison avec la souche symbiotique d’origine. CR Acad Sci Paris D. 1973;277: 981–984.

26. Stenbak CR, Ryu J-H, Leulier F, Pili-Floury S, Parquet C, Hervé M, et al. Peptidoglycan Molecular Requirements Allowing Detection by the Drosophila Immune Deficiency Pathway. J Immunol. 2004;173: 7339–7348. doi:10.4049/jimmunol.173.12.7339

27. Martin M. Cutadapt removes adapter sequences from high-throughput sequencing reads. EMBnet.journal. 2011;17: 10–12. doi:10.14806/ej.17.1.200

28. Dobin A, Davis CA, Schlesinger F, Drenkow J, Zaleski C, Jha S, et al. STAR: ultrafast universal RNA-seq aligner. Bioinformatics. 2013;29: 15–21. doi:10.1093/bioinformatics/bts635

29. Langmead B, Salzberg SL. Fast gapped-read alignment with Bowtie 2. Nat Methods. 2012;9: 357–359. doi:10.1038/nmeth.1923

30. Li H, Handsaker B, Wysoker A, Fennell T, Ruan J, Homer N, et al. The Sequence Alignment/Map format and SAMtools. Bioinformatics. 2009;25: 2078–2079. doi:10.1093/bioinformatics/btp352

31. Liao Y, Smyth GK, Shi W. The Subread aligner: fast, accurate and scalable read mapping by seed-and-vote. Nucleic Acids Res. 2013;41: e108. doi:10.1093/nar/gkt214

32. Lerat E, Fablet M, Modolo L, Lopez-Maestre H, Vieira C. TEtools facilitates big data expression analysis of transposable elements and reveals an antagonism between their activity and that of piRNA genes. Nucleic Acids Res. 2017;45: e17. doi:10.1093/nar/gkw953

33. Love MI, Huber W, Anders S. Moderated estimation of fold change and dispersion for RNA-seq data with DESeq2. Genome Biol. 2014;15: 550. doi:10.1186/s13059-014-0550-8

34. Benjamini Y, Hochberg Y. Controlling the False Discovery Rate: A Practical and Powerful Approach to Multiple Testing. J R Stat Soc Ser B Methodol. 1995;57: 289–300. doi:10.1111/j.2517-6161.1995.tb02031.x

35. Parisot N, Vargas-Chávez C, Goubert C, Baa-Puyoulet P, Balmand S, Beranger L, et al. The transposable element-rich genome of the cereal pest Sitophilus oryzae. BMC Biol. 2021;19: 241. doi:10.1186/s12915-021-01158-2

36. Ji J, Zhou L, Xu Z, Ma L, Lu Z. Two atypical gram-negative bacteria-binding proteins are involved in the antibacterial response in the pea aphid (Acyrthosiphon pisum). Insect Mol Biol. 2021;30: 427–435. doi:10.1111/imb.12708

37. Hughes AL. Evolution of the βGRP/GNBP/β-1,3-glucanase family of insects. Immunogenetics. 2012;64: 549–558. doi:10.1007/s00251-012-0610-8

38. Yamazaki Y, Matsunaga Y, Tokunaga Y, Obayashi S, Saito M, Morita T. Snake venom Vascular Endothelial Growth Factors (VEGF-Fs) exclusively vary their structures and functions among species. J Biol Chem. 2009;284: 9885–9891. doi:10.1074/jbc.M809071200

39. Sodani K, Patel A, Kathawala RJ, Chen Z-S. Multidrug resistance associated proteins in multidrug resistance. Chin J Cancer. 2012;31: 58–72. doi:10.5732/cjc.011.10329

40. Gottesman S. Trouble is coming: Signaling pathways that regulate general stress responses in bacteria. J Biol Chem. 2019;294: 11685–11700. doi:10.1074/jbc.REV119.005593

41. Ishihama A. Functional modulation of Escherichia coli RNA polymerase. Annu Rev Microbiol. 2000;54: 499–518. doi:10.1146/annurev.micro.54.1.499

42. Costechareyre D, Chich J-F, Strub J-M, Rahbé Y, Condemine G. Transcriptome of Dickeya dadantii Infecting Acyrthosiphon pisum Reveals a Strong Defense against Antimicrobial Peptides. PLOS ONE. 2013;8: e54118. doi:10.1371/journal.pone.0054118

43. Charles H, Heddi A, Guillaud J, Nardon C, Nardon P. A Molecular Aspect of Symbiotic Interactions between the WeevilSitophilus oryzaeand Its Endosymbiotic Bacteria: Over-expression of a Chaperonin. Biochem Biophys Res Commun. 1997;239: 769–774. doi:10.1006/bbrc.1997.7552

44. Kupper M, Gupta SK, Feldhaar H, Gross R. Versatile roles of the chaperonin GroEL in microorganism–insect interactions. FEMS Microbiol Lett. 2014;353: 1–10. doi:10.1111/1574-6968.12390

45. Fares MA, Moya A, Barrio E. GroEL and the maintenance of bacterial endosymbiosis. Trends Genet. 2004;20: 413–416. doi:10.1016/j.tig.2004.07.001

46. Fares MA, Ruiz-González MX, Moya A, Elena SF, Barrio E. GroEL buffers against deleterious mutations. Nature. 2002;417: 398–398. doi:10.1038/417398a

47. Meier EL, Goley ED. Form and function of the bacterial cytokinetic ring. Curr Opin Cell Biol. 2014;26: 19–27. doi:10.1016/j.ceb.2013.08.006

48. Eraso JM, Markillie LM, Mitchell HD, Taylor RC, Orr G, Margolin W. The Highly Conserved MraZ Protein Is a Transcriptional Regulator in Escherichia coli. J Bacteriol. 2014;196: 2053–2066. doi:10.1128/JB.01370-13

49. Pan J, Zhao M, Huang Y, Li J, Liu X, Ren Z, et al. Integration Host Factor Modulates the Expression and Function of T6SS2 in Vibrio fluvialis. Front Microbiol. 2018;9. doi:10.3389/fmicb.2018.00962

50. Sevin EW, Barloy-Hubler F. RASTA-Bacteria: a web-based tool for identifying toxin-antitoxin loci in prokaryotes. Genome Biol. 2007;8: R155. doi:10.1186/gb-2007-8-8-r155

51. Szekeres S, Dauti M, Wilde C, Mazel D, Rowe-Magnus DA. Chromosomal toxin–antitoxin loci can diminish large-scale genome reductions in the absence of selection. Mol Microbiol. 2007;63: 1588–1605. doi:10.1111/j.1365-2958.2007.05613.x

52. Manniello MD, Moretta A, Salvia R, Scieuzo C, Lucchetti D, Vogel H, et al. Insect antimicrobial peptides: potential weapons to counteract the antibiotic resistance. Cell Mol Life Sci. 2021;78: 4259–4282. doi:10.1007/s00018-021-03784-z

53. Maltz MA, Weiss BL, O’Neill M, Wu Y, Aksoy S. OmpA-Mediated Biofilm Formation Is Essential for the Commensal Bacterium Sodalis glossinidius To Colonize the Tsetse Fly Gut. Appl Environ Microbiol. 2012;78: 7760–7768. doi:10.1128/AEM.01858-12

54. Gerardo NM, Hoang KL, Stoy KS. Evolution of animal immunity in the light of beneficial symbioses. Philos Trans R Soc B Biol Sci. 2020;375: 20190601. doi:10.1098/rstb.2019.0601

55. Buchner P, Mueller B. Endosymbiosis of Animals with Plant Microorganisms. Wiley; 1965.

56. Wang D, Liu Y, Su Y, Wei C. Bacterial Communities in Bacteriomes, Ovaries and Testes of three Geographical Populations of a Sap-Feeding Insect, Platypleura kaempferi (Hemiptera: Cicadidae). Curr Microbiol. 2021;78: 1778–1791. doi:10.1007/s00284-021-02435-7

57. Wang D, Huang Z, Billen J, Zhang G, He H, Wei C. Structural diversity of symbionts and related cellular mechanisms underlying vertical symbiont transmission in cicadas. Environ Microbiol. 2021;23: 6603–6621. doi:10.1111/1462-2920.15711

58. Kucuk RA. Gut Bacteria in the Holometabola: A Review of Obligate and Facultative Symbionts. J Insect Sci Online. 2020;20: 22. doi:10.1093/jisesa/ieaa084

