## Additional text 1 for "Efficient compartmentalization in insect bacteriomes protects symbiotic bacteria from host immune system"

Analysis of the complete bacterial transcriptome from both controls and TCT-challenged larvae display similar gene expression, including within metabolic pathways, highlighting an active bacterial metabolism, which is also known to accelerate larval development and increase nutrient availability for the host [3]. Highly expressed bacterial protein coding genes within the bacteriome (Transcripts Per Million - TPM > 1000, Additional Table 8) were mainly involved in transcriptional regulation, translation, stress response, and virulence (Additional Table 9). Among the highest expressed genes we detected cold and heat shock protein coding genes (*cspA, cspE, rpoH*) and several chaperones (*groEL1/2*, *groES1/2*, *hlpA*). These results agree with previous studies that have detected the chaperonin GroEL as the most expressed protein in *S. pierantonius,* accounting for 40% of the bacterial protein synthesis [43]. It has been documented that the constitutive expression of the chaperonin GroEL (and possibly of other stress-response proteins) in endosymbionts with reduced genomes is essential to mitigate the deleterious effects of genome erosion, by assisting the folding of conformationally damaged proteins [44–46]. In weevils, GroEL was proposed to have a central role in the inhibition of *S. pierantonius* division, through interaction with ColA [18]. Other highly expressed genes related to cell division that could be involved in this inhibition are the cell division factor *zapA*, which localizes to the cytokinetic ring [47], and the cell division gene and transcriptional regulator *mraZ* [48]. It is interesting to note that three genes belonging to bacterial secretion systems (*secG*, *ssaD* and *invF*) were highly expressed in L4 larvae, suggesting the importance of these free-living bacterial infectious traits to this endosymbiont, possibly conferring additional virulence to this bacterium. Moreover, several transcriptional, translational and stabilization factors of the general stress response sigma factor RpoS (reviewed in [40]) were expressed at varied levels (Additional Table 8). The expression of *rpoS* in all conditions tested was lower than the vegetative sigma factor *rpoD*, which is a typical profile of the exponential growth phase in *Escherichia coli* [41]. This basal level of *rpoS* is also needed for triggering a fast stress response in diverse bacteria [40], and shows the ability of *S. pierantonius* from larval bacteriomes to quickly enter a "virulent mode" in the subsequent pupal stage in order to exit bacteriocytes and re-infect stem cells [16]. In addition, the gene *ihfA*, coding for the Integration Host Factor subunit Alpha, a bacterial protein that confers the propagation of antibiotic resistance and virulence factors in bacterial populations, was highly expressed and has been proposed to modulate the expression and function of type IV secretion system in the bacterial pathogen *Vibrio fluvialis* [49]. Moreover, when we compared highly expressed genes with differentially expressed genes throughout the metamorphosis of *S. oryzae* [16], we found that more than 50% were common to both groups of genes, which highlights the fact that not only these are important genes in terms of expression levels, but that *S. pierantonius* has the ability to modulate them throughout the weevil's life cycle. For instance, components of the T3SS were detected as up-regulated in pupae, a developmental stage in which endosymbionts are thought to enter a “virulent mode” and exit the bacteriocytes to re-infect stem cells [16]. Finally, we detected the expression of the chromosomal *hok*/*sok* toxin-antitoxin (TA) system. TA systems are essential regulators of growth arrest and programmed cell death which are found ubiquitously in free-living bacteria [50]. These systems were already proposed to diminish the deleterious effects of genome reduction in the absence of natural selection [51], and *S. pierantonius* has already been proposed as an interesting model organism for the study of endosymbiont genome reduction [15].

Overall, *S. pierantonius* seems to be in a dynamic equilibrium between fighting the deleterious effects of genome erosion occurring at initial steps of endosymbiosis, while maintaining the main mechanisms necessary for survival.
