## Supplementary figures and images for "Efficient compartmentalization in insect bacteriomes protects symbiotic bacteria from host immune system"

### Additional figure 1

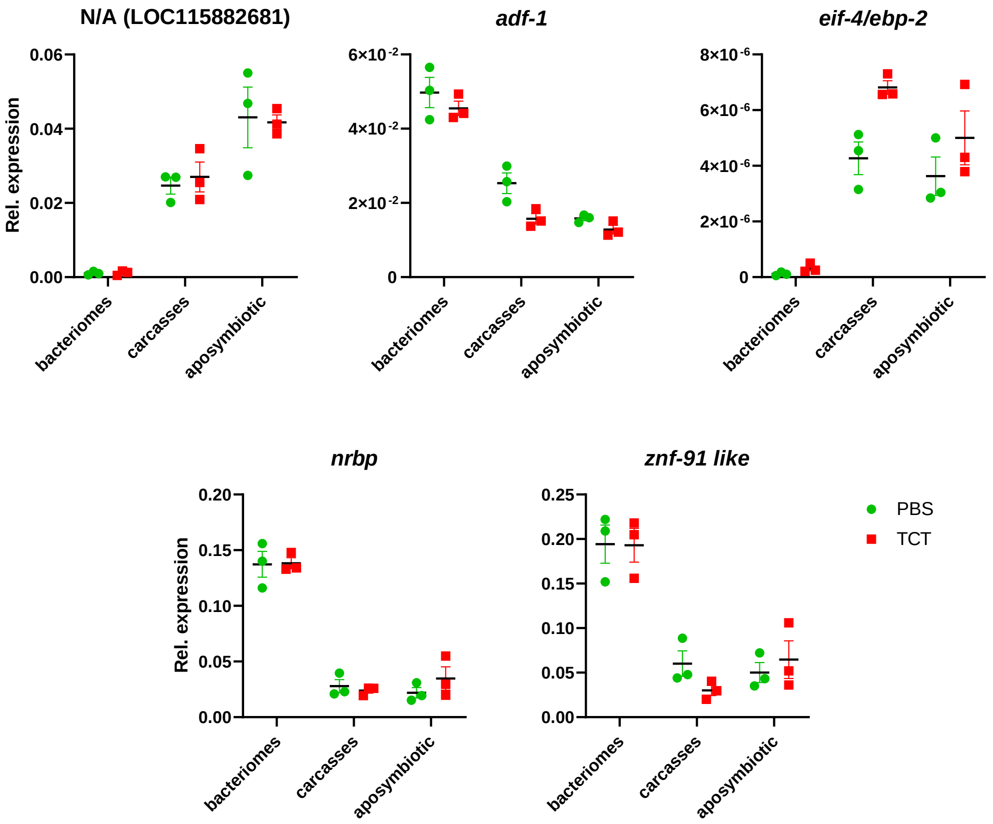
